# Whole genome sequence analyses of Western Central African Pygmy hunter-gatherers reveal a complex demographic history and identify candidate genes under positive natural selection

**DOI:** 10.1101/022194

**Authors:** PingHsun Hsieh, Krishna R. Veeramah, Joseph Lachance, Sarah A. Tishkoff, Jeffrey D. Wall, Michael F. Hammer, Ryan N. Gutenkunst

**Affiliations:** Department of Ecology and Evolutionary Biology, University of Arizona, Tucson, AZ; Arizona Research Laboratories Division of Biotechnology, University of Arizona, Tucson, AZ; Department of Molecular and Cellular Biology, University of Arizona, Tucson, AZ; Department of Biology and Genetics, University of Pennsylvania, Philadelphia, PA; Institute for Human Genetics, University of California, San Francisco, CA; Department of Ecology and Evolution, Stony Brook University, Stony Brook, NY; School of Biology, Georgia Institute of Technology, Atlanta, GA

**Keywords:** African Pygmies, demographic inference, natural selection, whole-genome scan

## Abstract

African Pygmies practicing a mobile hunter-gatherer lifestyle are phenotypically and genetically diverged from other anatomically modern humans, and they likely experienced strong selective pressures due to their unique lifestyle in the Central African rainforest. To identify genomic targets of adaptation, we sequenced the genomes of four Biaka Pygmies from the Central African Republic and jointly analyzed these data with the genome sequences of three Baka Pygmies from Cameroon and nine Yoruba famers. To account for the complex demographic history of these populations that includes both isolation and gene flow, we fit models using the joint allele frequency spectrum and validated them using independent approaches. Our two best-fit models both suggest ancient divergence between the ancestors of the farmers and Pygmies, 90,000 or 150,000 years ago. We also find that bi-directional asymmetric gene-flow is statistically better supported than a single pulse of unidirectional gene flow from farmers to Pygmies, as previously suggested. We then applied complementary statistics to scan the genome for evidence of selective sweeps and polygenic selection. We found that conventional statistical outlier approaches were biased toward identifying candidates in regions of high mutation or low recombination rate. To avoid this bias, we assigned *P*-values for candidates using whole-genome simulations incorporating demography and variation in both recombination and mutation rates. We found that genes and gene sets involved in muscle development, bone synthesis, immunity, reproduction, cell signaling and development, and energy metabolism are likely to be targets of positive natural selection in Western African Pygmies or their recent ancestors.

## Introduction

Recent archaeological and genetic studies suggest that anatomically modern humans (AMH) originated in Africa prior to 160-190 thousand year ago (kya, Cavalli-Sforza 1986). Before the invention of agriculture in the Neolithic (~6–10 kya), hunting and gathering was the subsistence strategy employed by early human societies (Cavalli-Sforza 1986; Henshilwood et al. 2002; Scheinfeldt et al. 2010). Among extant African human populations, the Pygmies, commonly identified by their short stature (mean adult height < 150–160 cm), are one of the few populations that still predominantly practice a hunting and gathering lifestyle. The common view in archeology is that the Pygmy people have been forest dwellers, dating from at least 40 kya (Cavalli-Sforza 1986).

African Pygmies forage in the equatorial rainforests of Central Africa and can be divided into Western and Eastern Pygmies. Western Pygmies (e.g., Baka and Biaka) mainly reside in rainforest west of the Congo Basin, while Eastern Pygmies (e.g., Mbuti and Efe) live in and around the Ituri rainforest and further south extending toward Lake Victoria (Cavalli-Sforza et al. 1994). Although still living as mobile hunter-gatherers, moving from one camping site to another in the forest regularly, archeological and cultural studies indicate that Pygmies have established social and economic contacts with nearby settled farmers (Cavalli-Sforza et al. 1994). For example, in addition to hunting animals and collecting plant foods, such as yams and honey, the Efe Pygmies trade forest food to Lese farmers in exchange for cultivated goods (Terashima 1987; Terashima 1998). Moreover, most Pygmies now speak Niger-Kordofanian (e.g. Bantu) or Nilo-Saharan languages, possibly acquired from neighboring farmers, especially since the expansion of Bantu-speaking agriculturalists, beginning about 5 kya (Blench 2006).

Recent genetic evidence favors a single origin of African Pygmies (Patin et al. 2009; Batini et al. 2011; Veeramah et al. 2011).Western Pygmies have likely experienced greater genetic admixture with neighboring non-Pygmy farmer populations than Eastern Pygmies (Cavalli-Sforza 1986; Patin et al. 2009; Tishkoff et al. 2009; Veeramah et al. 2011; Verdu et al. 2013). Several mitochondrial and multi-locus DNA studies have estimated that African Pygmies diverged from the ancestors of present-day Niger-Cordofanian agriculturalists ~60 kya (95% C.I.: 25–130 kya, Patin et al. 2009), ~70 kya (95% C.I.: 51– 106 kya, Batini et al. 2011), and ~49 kya (95% C.I.: 10-105 kya, Veeramah et al. 2011). However, because each of these studies employed less than 60 genomic loci, these studies either made strong *a priori* assumptions to restrict the parameter space they searched in their demographic modeling (Patin et al. 2009) or did not have sufficient statistical power to infer gene flow between populations (Batini et al. 2011, Veeramah et al. 2011). Thus, a comprehensive understanding of the demographic prehistory of African Pygmies remains lacking.

Pygmy populations have long been studied biologically and genetically because of their distinct phenotypes, particularly short stature. Physiological evidence suggests that short stature is associated with low growth hormone binding protein and insulin-like growth factor-I (IGF-I) levels in Pygmy groups in different parts of the world (Baumann et al. 1989; Dávila et al. 2002). Using high-density SNP chip data, several population genetic studies have reported candidates for Pygmy short stature, including genes in the IGF-I pathway (Pickrell et al. 2009; Jarvis et al. 2012; Migliano et al. 2013), the iodine-dependent thyroid hormone pathway (Herráez et al. 2009; Migliano et al. 2013), and the bone homeostatsis/skeletal remodeling pathway (Mendizabal et al. 2012). Lachance et al. (2012) searched signals of positive selection in five high-coverage Western Pygmy genomes and suggested that short stature may be due to selection on genes involved in development of the anterior pituitary, as well as the crosstalk between the adiponectin and insulin-singling pathways. A more recent study using admixture mapping identified 16 regions associated with height in Batwa Pygmies, which were enriched for SNPs associated with height in Europeans and for genes with growth hormone receptor and regulation functions (Perry et al. 2014).

Several hypotheses have been proposed regarding Pygmy adaptation to the dense, humid forest environment, all of which may influence stature. These include thermoregulatory adaptation to the tropical forest (Cavalli-Sforza 1986), reduction of caloric intake in a food-limited environment (Shea and Bailey 1996), improved mobility in the dense forest (Diamond 1991), and earlier reproduction to compensate for short lifespans (Migliano et al. 2007). In addition, the equatorial rainforest in Central Africa is enriched in pathogens and parasites, such as malaria and haemorrhagic fever (Ohenjo et al. 2006). Loci that are involved in immunity have thus been suggested to be targets for adaptation to this challenging forest environment (Jarvis et al. 2012; Lachance et al. 2012).

Although previous studies have identified many possible targets of adaptive selection in African Pygmies, challenges remain. First, demographic events and local genomic architecture (e.g., heterogeneity in mutation and recombination rates) can mimic the genetic patterns generated by adaptation (Schaffner et al. 2005; Teshima et al. 2006). High false positive and false negative rates are expected in studies that determine candidates of natural selection based solely on selecting outliers from the distribution of a test statistic (Jeffreys et al. 2005; Schaffner et al. 2005; Teshima et al. 2006; Akey 2009). In addition, the large genomic sizes of candidate regions (on the order of 100 kb), especially for those reported in SNP-microarray studies, make inference of the genetic basis of adaptation difficult.

Understanding genetic adaptation in African Pygmies, therefore, requires not only leveraging high-coverage whole-genome data to perform a thorough scan for selective signatures, but also realistic demographic models to assess the statistical significance of the candidates. To provide a genomic perspective on adaptation in Pygmies, we sequenced four western Biaka Pygmies from the Central African Republic using Complete Genomics (CGI) technology (Drmanac et al. 2010) and combined these data with similar data from three Baka Pygmies (Lachance et al. 2012) from Cameroon and nine unrelated Yoruba farmers. We inferred the demographic history of these populations and searched for positive selection using several complementary statistical methods. We assessed statistical significance in our selection scan analyses using genome-scale simulations performed with MaCS (Chen et al. 2009) that incorporated recombination and mutation rate heterogeneity along the genome. Finally, we functionally annotated our candidates, and we discuss their biological impact. Our analysis thus provides unique insights into the complex demographic and adaptive history of Western African Pygmies.

## Results

### Demographic history inference for West African Pygmies and Farmers

We used the demographic inference tool *∂*a*∂*i (Gutenkunst et al. 2009) to infer the joint demographic history of one farmer (Yoruba) and two Pygmy (Baka and Biaka) populations using our high coverage (median=60.5X) CGI whole genome data. After removing single nucleotide variant (SNVs) that failed our quality control criteria (see Materials and Methods), we used 1.58 million intergenic autosomal SNVs to build a 3-population unfolded allele frequency spectrum (AFS). Ancestral states were inferred using chimpanzee as the outgroup. The estimated sequence divergence between human and chimpanzee based on these non-genic sequences is 1.14%. Because misspecification of ancestral states might alter the AFS and lead to patterns that can mimic positive selection through an excess of high frequency variants, we used the method of Hernandez et al. (2007) implemented in *∂*a*∂*i to statistically correct the AFS for the ancestral misspecification. Because linkage among sites means that *∂*a*∂*i calculates a composite likelihood rather than the full likelihood, we estimated confidence intervals via conventional non-parametric bootstraps (see Materials and Methods).

To guide development of three-population models, we first considered simpler one- and two-population models. These initial simpler models consistently suggested a more recent divergence between the two Pygmy populations than between either of those populations and the farmers. Based on these results and previously published inferences (Patin et al. 2009; Batini et al. 2011; Veeramah et al. 2011; Verdu et al. 2013), we tested multiple three-population models, considering a variety of scenarios for gene flow and population size changes. The best-fit three-population demographic model, Model-1 (continuous asymmetric gene flow, composite log-likelihood= –6,712), is illustrated in **Figure 1A**. The maximum composite-likelihood estimates for the 10 free parameters are reported in **Table 1**. The joint frequency spectra resulting from this model qualitatively reproduce the data, as seen in the second and third rows of **Figure 1C**, although our model does produce an excess of high frequency shared variants. In Model-1, the ancestors of contemporary farmers and Pygmies diverged ~156 kya (95% C.I.: 140–164 kya) from an ancestral population that had expanded roughly three-fold prior to divergence. The ancestors of the farmers and Pygmies remained isolated until ~40 kya (95% C.I.: 36–44 kya), at which point bi-directional gene flow began, with the flow from farmers to Pygmies being 10 times greater than from Pygmies to farmers (**Table 1**). Following the Pygmy-farmer divergence, the effective population size of farmers increased and the effective population size of Pygmies decreased. The Baka and Biaka diverged much more recently, about 5 kya (95% C.I.: 4.7–5.7 kya). Because our small sample size limits the power to infer recent demographic events (Robinson et al. 2014), we assumed that the Baka-Biaka divergence did not change the rates of gene flow with the Yoruba, and our model includes no Baka-Biaka gene flow.

**Figure 1.**
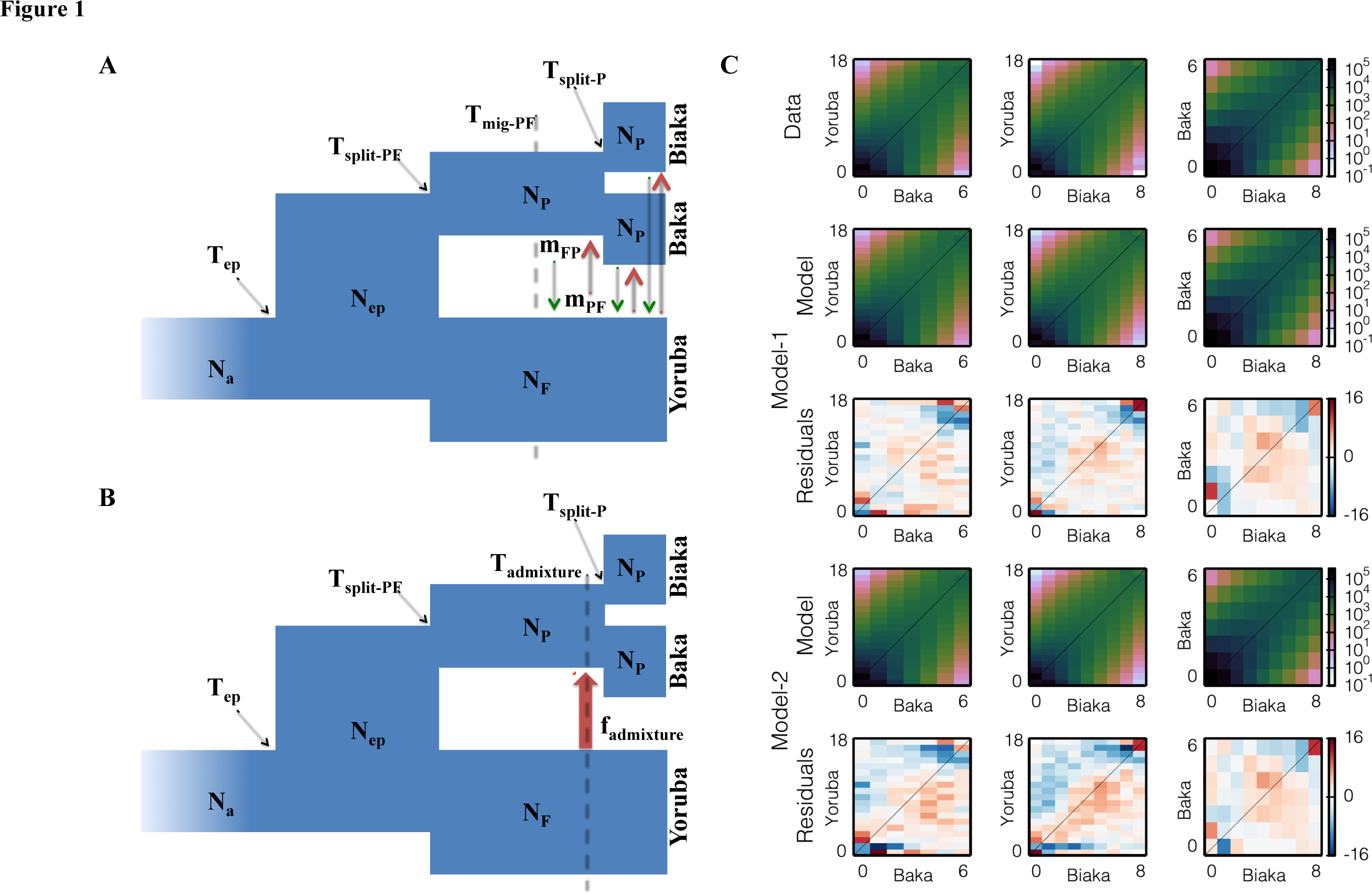
Best-fit demographic models and observed and predicted frequency spectra for African farmer (Yoruba) and Pygmy (Baka and Biaka) populations. (A) The continuous asymmetric gene flow model (Model-1) we fit, with the 10 free parameters labeled. (B) The single-pulse admixture model (Model-2) we fit, with the 9 free parameters labeled. (C) The marginal spectra for each pair of populations. Row one is data, row two (Model-1) and four (Model-2) are models, and row three and five are Anscombe residuals of model minus data for Model-1 and Model-2, respectively.

**Table 1.** Parameter estimates and confidence intervals for two best-fit demographic models. Model-1: continuous asymmetric gene flow. Model-: single-pulse gene flow. Estimates and confidence intervals are shown for effective population sizes (N), times (T) of population divergence and gene low onset, levels of gene flow (m) between farmer (F) and Pygmy (P) populations. T_admixture_ and f_admixture_ refer to the timing and strength of the single-pulse gene flow from the farmers (F) to Pygmies (P) in Model-2.

| Demographic parameters | Model-1<br>(Asymmetric gene flow) |  | Model-2<br>(Single-pulse gene flow) |  |
| --- | --- | --- | --- | --- |
|  | Estimates | <sup>†</sup>95% C.I. | Estimates | <sup>†</sup>95% C.I. |
| $N_a$: $N_e$ ancestral population | 6,727 | 6,676 – 6,819 | 6,735 | 6,671– 6,826 |
| $N_{ep}$: $N_e$ ancestral population after expansion | 20,473 | 15,560 – 27,561 | 15,236 | 14,436 – 15,894 |
| $N_F$: $N_e$ contemporary Farmer (F) | 11,900 | 11,714 – 12,138 | 13,854 | 13,721 – 14,055 |
| $N_P$: $N_e$ contemporary Pygmy (P) | 5,831 | 5,631 – 5,986 | 5,373 | 5,217 – 5,530 |
| $T_{ep}$: Time<sup>†</sup> of ancestral expansion | 221,118 | 210,513 – 236,634 | 232,629 | 223,172 – 244,327 |
| $T_{\text{split-PF}}$: Time of P-F split | 155,671 | 139,661 – 164,280 | 89,645 | 85,503– 91,725 |
| $T_{\text{mig-PF}}$: Time of onset of gene flow between P and F | 39,337 | 36,565 – 43,550 | – | – |
| $T_{\text{admixture}}$: Time of admixture from F to P | – | – | 7,136 | 6,887 – 7,656 |
| $T_{\text{split-P}}$: Time of split between the two P populations | 5,139 | 4,762 – 5,630 | 4,049 | 3,803 - 4,396 |
| $m_{PF}$: Gene flow<sup>‡</sup> ($P \leftarrow F$) | $9.0 \times 10^{-4}$ | $8.4 \times 10^{-4}$ – $9.4 \times 10^{-4}$ | – | – |
| $m_{FP}$: Gene flow ($F \leftarrow P$) | $9.1 \times 10^{-5}$ | $8.2 \times 10^{-5}$ - $1 \times 10^{-4}$ | – | – |
| $f_{\text{admixture}}$: Strength of admixture ($P \leftarrow F$) | – | – | 0.6799 | 0.6789 – 0.6818 |

<sup>\*</sup>Effective population size in individuals. <sup>†</sup>Time in years, assuming 25 years per generation and mutation rate $2.35 \times 10^{-8}$ per base per generation (Gutenkunst et al. 2009). <sup>‡</sup>Fraction of the population each generation that are new migrants. <sup>†</sup>Confidence intervals estimated using 100 conventional bootstraps.

Our second best-fit model involves a recent pulse of unidirectional gene flow from farmers to Pygmies (Model-2, **Figure 1B and 1C**) after the divergence of the two populations. The maximum composite-likelihood estimates for the 9 free parameters are shown in Table 1. The maximum composite log-likelihood of Model-2 (-7,737) is lower than Model-1. In Model-2, we inferred that Pygmies and farmers diverged about 90 kya (95% C.I.: 85–92 kya). The pulse of gene flow is estimated to have occurred ~7 kya (95% C.I.: 6.8–7.7 kya), while the inferred admixture proportion in our Pygmy sample resulting from the pulse of gene flow from the farmers is ~68% (C.I.: 67.9–68.2%).

### Model Selection and Validation of Demographic inference

We used three approaches to validate our demographic inference (see Materials and Methods). First, to remove the effects of linkage we refit our models to a subset of the data in which variant sites were at least 0.01 centiMorgan (cM) apart. The two best-fitting models remained the same as using the whole dataset, and the parameter estimates were compatible (**Table S1**). Under the assumption that the likelihoods calculated using the thinned dataset are full likelihoods, we applied the Akaike (AIC, Akaike 1974) and Bayesian information criteria (BIC, Schwarz 1978) for model selection. Both AIC and BIC prefer the continuous asymmetric gene-flow model to the single-pulse gene flow model (**Table S1**).

As a second validation, we used patterns of linkage disequilibrium (LD) decay, information not utilized by *∂*a*∂*i. We calculated LD using sliding windows of 0.1 cM in the real data and in simulated whole-genome data, using 100 models drawn from the parameter confidence intervals of our two best-fit demographic models. We found that the patterns of LD decay predicted by the models generally matched the data well for both Pygmies and farmers (**Figure S1**). In Pygmies, Model-1 matches the LD decay in the real data for pairs of sites close to or far from each other, but not for sites with intermediate separation (**Figure S1A**). On the other hand, Model-2 matches the data for pairs of sites with intermediate and large separations, but not for small separations. Similar differences are observed for the farmers (**Figure S1B**). These discrepancies suggest that neither of our two best-fit models perfectly captures the full demographic history of our populations, although they do capture important features of that history.

As a third validation, we applied the pairwise sequentially Markovian coalescent (PSMC, Li and Durbin 2011) as an independent means to explore the demographic history of our populations (Materials and Methods). PSMC infers effective population size over time from a single diploid genome. We applied PSMC to our whole genome non-genic data that pass our quality control metrics. The PSMC curves of the farmers begin to separate from those of the Pygmies roughly 100–200 kya and are completely separated by about 70–90 kya (**Figure S2**). The separation of PSMC curves of individuals from different populations is evidence of population divergence (Li and Durbin 2011). Thus, our PSMC analysis suggests that the ancestors of the farmers and Pygmies began differentiating from each other as early as 100–200 kya, consistent with the inferred divergence time in Model-1. To test whether Model-1 and/or Model-2 recapitulates the observed deep divergence time between farmers and Pygmies seen via PSMC, we applied PSMC to two simulated genomes under both models for the farmer and Pygmy groups, respectively (**Figure S3**). Under Model-1, the PSMC curves of the simulated Pygmy genomes depart from those of the simulated farmer genomes at about the same time as in the PSMC analysis of the real data (**Figure S3A**), while the PSMC curves of the two groups simulated under Model-2 does not show clear separation until ~70 kya (**Figure S3B**). Together, these results suggest that that the ancient divergence time inferred using Model-1 is plausible.

In general, these validations suggest that Model-1 is our best estimate of demographic history for these populations, but it is an imperfect model. In order to lessen the impact of model misspecification on our selection inference, we conservatively report candidates under both Model-1 and Model-2.

### Prioritizing Selection Candidates using Whole Genome Demographic Simulations

Because conventional statistical outlier approaches are prone to false positives, we used whole-genome simulations under our realistic demographic models to assign statistical significance (*P*-values) in our selection scan. We used the coalescent simulator MaCS (Chen et al. 2009) in order to model both recombination- and mutation-rate heterogeneity across the entire genome. To assess possible biases on selection inference due to imperfection of the genetic recombination map, we ran two sets of simulations, using two published recombination maps: the African American genetic recombination map (Hinch et al. 2011) and the HapMap Yoruba genetic recombination map (Frazer et al. 2007) (Materials and Methods).

Methods for detecting natural selection often rely on summaries of local genetic variation, and they may be biased by variation in mutation rate across the human genome (Reich et al. 2002; Drake et al. 2005; Schaffner et al. 2005; Sainudiin et al. 2007). For example, G2D values (Nielsen et al. 2009) are correlated with local genetic diversity (Pearson correlation 0.298, p<2.2×10^-16^, **Figure S4**). We addressed this by estimating and incorporating local mutation rate variation across the whole genome in our simulations (Materials and Methods). Our whole-genome neutral simulation approach reproduces the pattern of local genetic diversity in the real data well (Pearson correlation=0.902, **Figure S5**). To assess whether mutation-rate heterogeneity could bias downstream inferences of selection, we compared results using two different sets of simulations under Model-1 to assign *P*-values. In the first set, the local mutation rate for each window was assigned to be the mean rate of the recombination decile to which that window belonged (**Figure S6**). In the second set, we estimated a local mutation rate for each window individually (**Figure S7**). The *P*-value distributions of G2D based on these two sets of simulations were calculated, and for both analyses we chose the top 0.5% windows in the *P*-value distributions as the top-hits. There is a clear shift to larger heterozygosity (estimated using “/base) for the top hits in the first simulation set (**Figure S6A**), compared with the second set (**Figure S7A**). As expected, the top hits in the first simulation set tended to be windows with larger numbers of variants, while the top hits from the second set were distributed across the whole range of observed heterozygosity across the genome (**Figure S6B** vs. **Figure S7B**). This suggests that incorrectly incorporating mutation rate variation in whole-genome simulations might lead to biases toward regions with unusually high mutation rate as candidates of natural selection. From here on, we thus used the per-window mutation rate approach (Materials and Methods) for all of the simulations. We recognize that this approach may discount some selection signals, yielding a more conservative inference of natural selection.

To identify genomic regions with signals of selection, we used sliding windows of 500 consecutive SNVs that pass our quality control metrics, with a step size of 100 SNVs. We first assessed the statistical significance of each window using 1,000 neutral whole genome simulations with parameters drawn from the confidence intervals of each of the two best-fit demographic models. Our top hits are the top 0.5% of windows in the *P*-value distribution of each test statistic. For finer *P*-value resolution, for each of our top candidates we then performed an additional 100,000 local simulations.

The distribution of *P*-values was sensitive to the genetic recombination map used in the simulations (**Figure S8**). In particular, the distribution of G2D p-values using the African American recombination map (Hinch et al. 2011) is shifted more toward p=1 than using the Yoruba HapMap recombination map, suggesting that inference using the African American recombination map would be more conservative (**Figure S8**). To avoid potential biases due to the choice of map and/or null model, we restricted our list of candidates to those that are top hits using all four combinations of the two recombination maps and the two best-fit demographic models. Because the *P*-value distributions based on the two null demographic models are highly correlated (Pearson correlation=0.984, p<2.2×10^-16^, **Figure S9**), and the analysis based on the African American genetic recombination model is more conservative, unless mentioned otherwise we report the *P*-values and false discovery rates obtained using Model-1 and the African American recombination map for our candidates.

To illustrate the importance of using *P*-values to determine candidates, rather than relying on outliers in the distribution of a test statistic, we plotted the *P*-value based on Model-1 as a function of the G2D statistic for each of the windows that we surveyed (**Figure 2;** similar result holds for the iHS analysis, **Figure S10**). Quadrant I contains the many windows that have extreme G2D values but are not statistically significant when the confounding effects of demography and genomic architecture are controlled for. Conversely, Quadrant III contains the many windows that are statistically significant even though their G2D values are not extreme on a genome-wide basis. Because the association between functional elements (e.g. exon and regulatory sequences) and selection is not expected if a large fraction of significant tests are false positives, we validated our *P*-value approach by comparing the spatial distribution of our candidates for selection with the distribution of known functional sequences in the genome (Voight et al. 2006; Williamson et al. 2007; Mendizabal et al. 2012). As expected, we found that our top hits of the *P*-value approach were enriched in exons of genes (**Table S2**; one-sided Fisher exact test, p=0.029). Interestingly, we find no enrichment of top hits in regions deemed functional based on five types of ENCODE (Gerstein et al. 2012) regulatory elements (Materials and Methods; **Table S2**).

**Figure 2.**
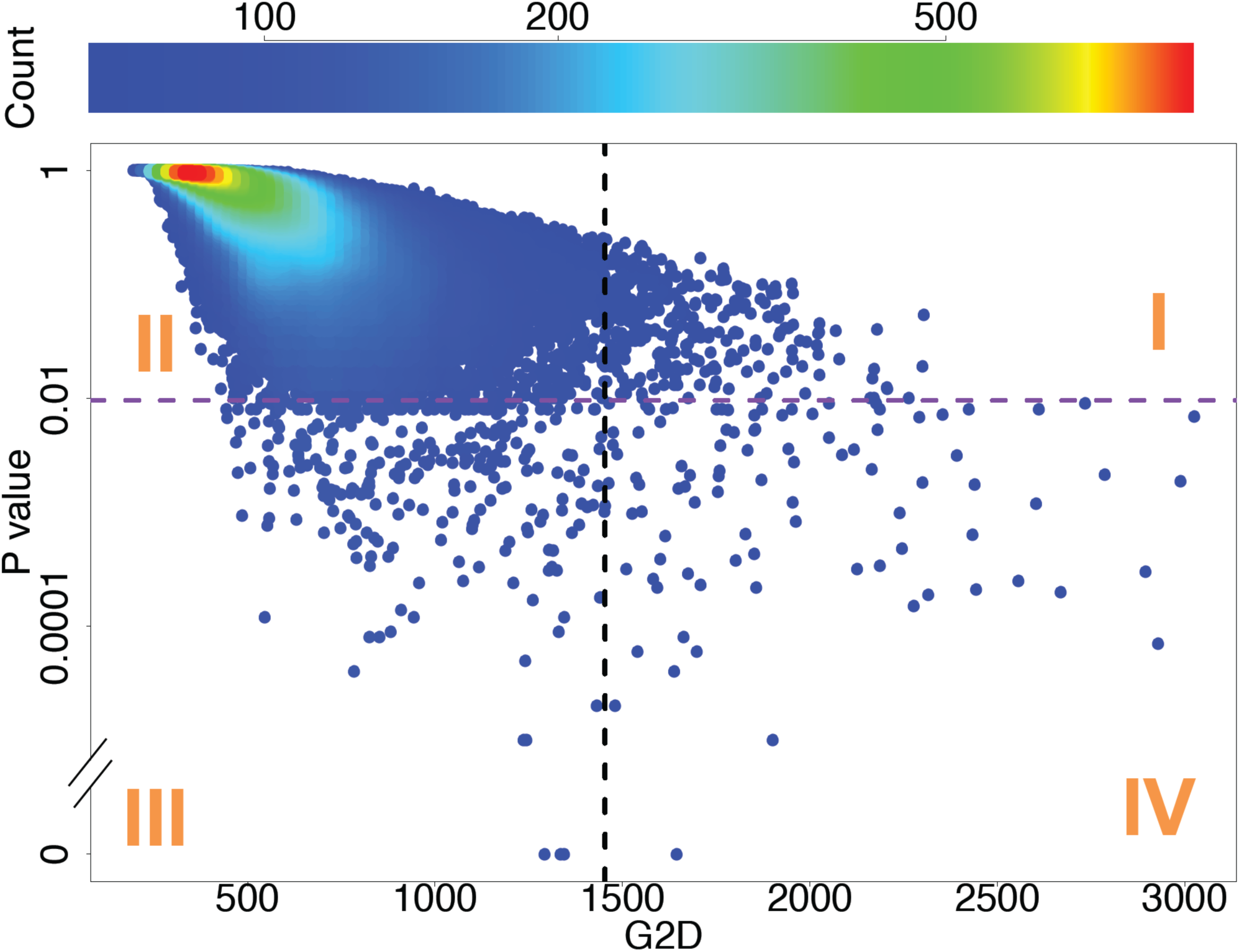
Importance of using *P*-values to define candidates in the G2D analysis. Each point is a window of 500 single nucleotide variants, and color represents the density of points. The vertical black line and the horizontal purple line are the top 0.5% significance cutoffs for the G2D and *P*-value distributions, respectively. Windows in Quadrant I are outliers in the G2D distribution but are not statistically significant when the effects of demography and genome architecture are controlled for. In Quadrant III are the many windows that are statistically significant even though their G2D values are modest.

### Evidence of Local Adaptation in Western African Pygmies: iHS

To detect recent incomplete selective sweeps, we scanned the genome using the haplotype-based iHS statistic (Voight et al. 2006) for the farmer and Pygmy samples separately. Each window was scored by the proportion of SNVs with standardized iHS score greater than 2, and the per-window *P*-value is the fraction of simulations in which that window’s score exceeded that in the real data (Materials and Methods). Using all four simulation sets, we defined Pygmy-specific signals as those windows that were a top-hit (the top 0.5% in the *P*-value distribution) in the Pygmy sample, but not in the Yoruba sample (not within the top 1% in the *P*-value distribution), yielding 35 distinct genomic regions (**Table S3**). We used a looser *P*-value cutoff to define Yoruba top-hits in order to be more conservative in identifying regions as Pygmy-specific.

Five of our candidate regions contain genes associated with bone synthesis. *EPHB1,* (locus: chr3:134572433-134716365, **Figure 3A**) is an *Ephrin* receptor at sites of osteogenesis. Interestingly, this region has been previously associated with the short stature in Pygmies (Jarvis et al. 2012). Our candidate region spans ~140 kb, containing exon 2 and exon 3 of *EPHB1* (which has a size of >460 kb and 16 coding exons). Elevated F_ST_ has been widely used to infer selection (Nielsen et al. 2009; Pickrell et al. 2009; Jarvis et al. 2012), and F_ST_ is elevated in this region, although we found no non-synonymous variants. To further investigate the signal of selection, we used hierarchical clustering and network analysis of the phased haplotypes for the region around exon 3 (± 10kb). Interestingly, both analyses suggest that Pygmy and farmer groups are almost fixed for different haplotypes (**Figure 3B-C**). This is consistent with an incomplete selective sweep (Voight et al. 2006; Pickrell et al. 2009; Pritchard et al. 2010) and indicative of different selective pressures in these two groups. The other four bone-synthesis related candidates are *SLCO2A1* (locus: chr3:133506737-133863702), *ZBTB38* (locus: chr3:141105569-141333249), *TSPAN5* (locus: chr4:99496207-99673561), and *GAREM* (locus: chr18:29766032-29896024). *SLCO2A1* encodes a prostaglandin transporter protein, and mutations in this gene have been shown causing Primary Hypertrophic Osteoarthropathy, a rare genetic disease that affects both skin and bones (Zhang et al. 2012). *ZBTB38* encodes a zinc finger transcriptional activator expressed in the brain, and has been associated with adult height in multiple populations (Lettre et al. 2008; Weedon et al. 2008; Wang et al. 2013). *TSPAN5* is a member of the tetraspanin protein family and is up-regulated during osteoclast differentiation (Iwai et al. 2007); knockdown of its expression dramatically inhibits osteoclastogenesis in vitro (Iwai et al. 2007; Zhou et al. 2014), suggesting its regulatory role in bone development. *GAREM* is an adapter protein in intracellular signaling cascades and has recently been associated with human height in a whole-exome sequencing association study (Kim et al. 2012). A few large F_ST_ (# 0.2) non-synonymous amino acid substitutions were observed within these candidate regions, but they are not suggested as functionally important by SIFT (Kumar et al. 2009) or PolyPhen2 (Adzhubei et al. 2010). Regions near four out of these five genes, however, show high levels of differentiated SNVs in enhancer and/or Polycomb-repressed sequences, implying that Pygmy short stature might arise partly through *cis*-regulatory evolution (**Figure S11**).

**Figure 3.**
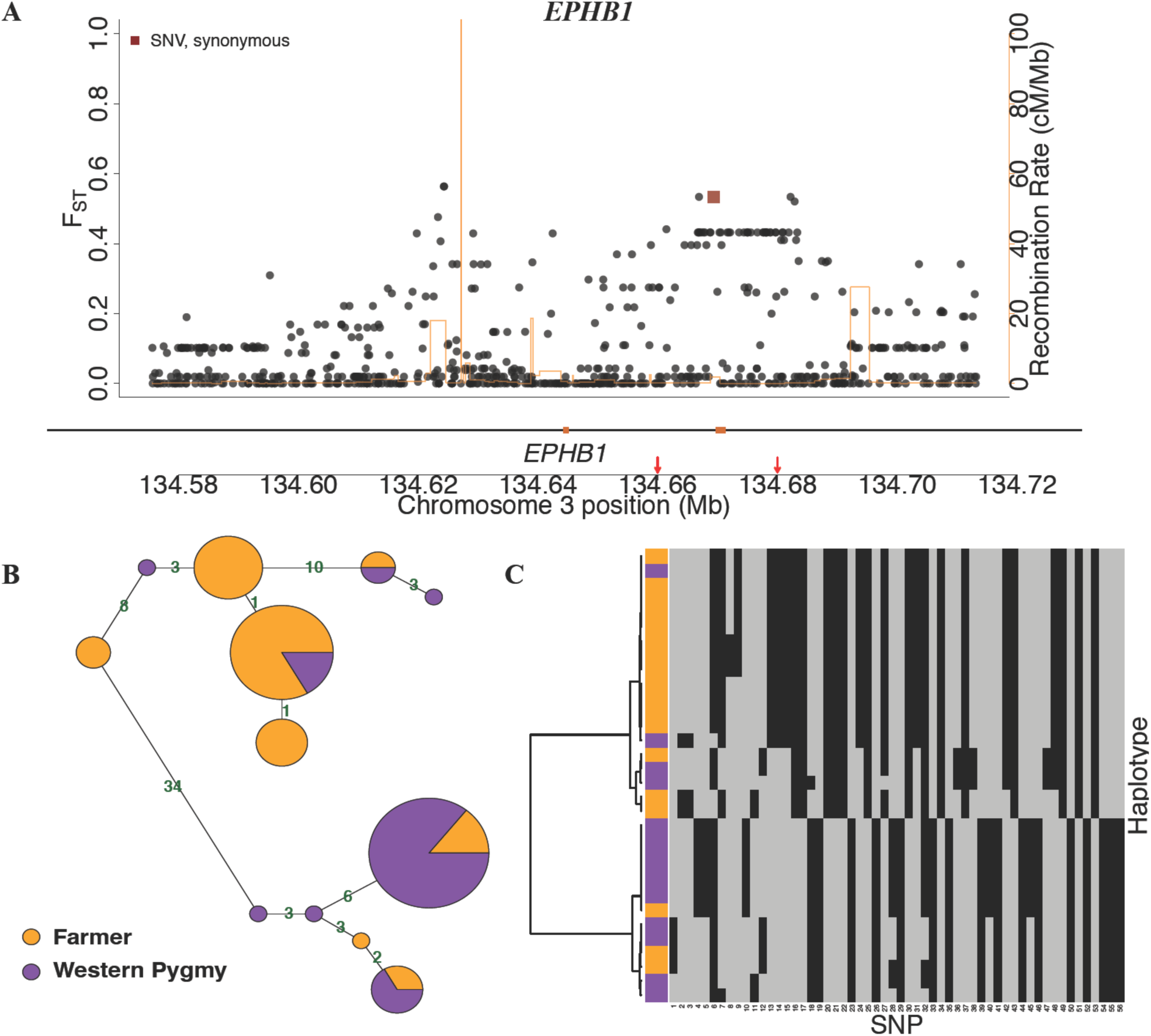
Candidate selection signal in *EPHB1*. (A) Distribution of F_ST_, functional annotation (Refseq and ENCODE elements, see Materials and Methods), and recombination rate (Hinch et al. 2011) for all variants (dots) in the candidate locus chr3:134572433-134716365. Genes are shown under the plot with black (non-coding sequences) and brown (exons) lines. (B) Haplotype network for the region (chr3: 134.66-134.68 Mb, the region between the two arrows in panel A) with elevated F_ST_. Each circle is a haplotype with size proportional to the number of chromosomes compared to the size of single chromosome shown on the legend and colors indicating the counts of that haplotype in our farmer and Pygmy samples. Haplotypes are connected by lines indicating the pairwise nucleotide distance between them. (C) Hierarchical clustering of the haplotypes in (B). Columns are SNPs, with grey and black for ancestral and derived alleles, respectively, while rows show individual haplotypes.

Consistent with the hypothesis of selection for mobility (Diamond 1991), we found candidate loci in several muscle-related genes. In particular, *OBSCN* (spans >150 kb with 81 exons within the candidate locus chr1:228103665-228842760, **Figure 4A**), an obscurin gene, has an important role in the organization of myofibrils during assembly and may mediate interactions between the sarcoplasmic reticulum (striated muscle fibers found in the skeletal system) and myofibrils (Young et al. 2001; Ackermann et al. 2014). Within this gene, 16 out of 46 non-synonymous amino acid variants are predicted as functionally important by either SIFT or PolyPhen2. The SNV with the largest F_ST_ (chr1:228475848, rs437129, F_ST_ = 0.54) in this region is fixed for the ancestral allele (Guanine, PanTro3, Hg19) in our Pygmy sample but is segregating at much lower frequency in our Yoruba farmer sample (allele frequency for G = 0.39 or 7/18), in both homozygote and heterozygote forms. The ancestral allele (G) frequencies of rs437129 in Yoruba, Luhya, and African American based on the 1000 Genome Project (Phase I) are 0.551, 0.665, and 0.590 (dbSNP 137). Analyses of the haplotypes between the two non-synonymous sites with F_ST_ > 0.5 (chr1:228475848 and chr1:228520973, including the 10kb flanking region; **Figure 4B-C**) suggest the existence of two major haplotypes in our sample that are relatively population-specific. We thus postulate that natural selection might have acted in different directions for this region between these two groups. Other muscle-related genes identified by our scan include *COX10* (locus: chr17:13911228-14241158) and *LARGE* (locus: chr22:34224706-34359718). *COX10* is a cytochrome c oxidase, and Diaz et al. (2005) reported that *COX10* knockout mice develop a slowly progressive myopathy. *LARGE* is a member of the N-acetylglucosaminyltransferase gene family, and mutations in this gene cause a form of congenital muscular dystrophy (Longman et al. 2003).

**Figure 4.**
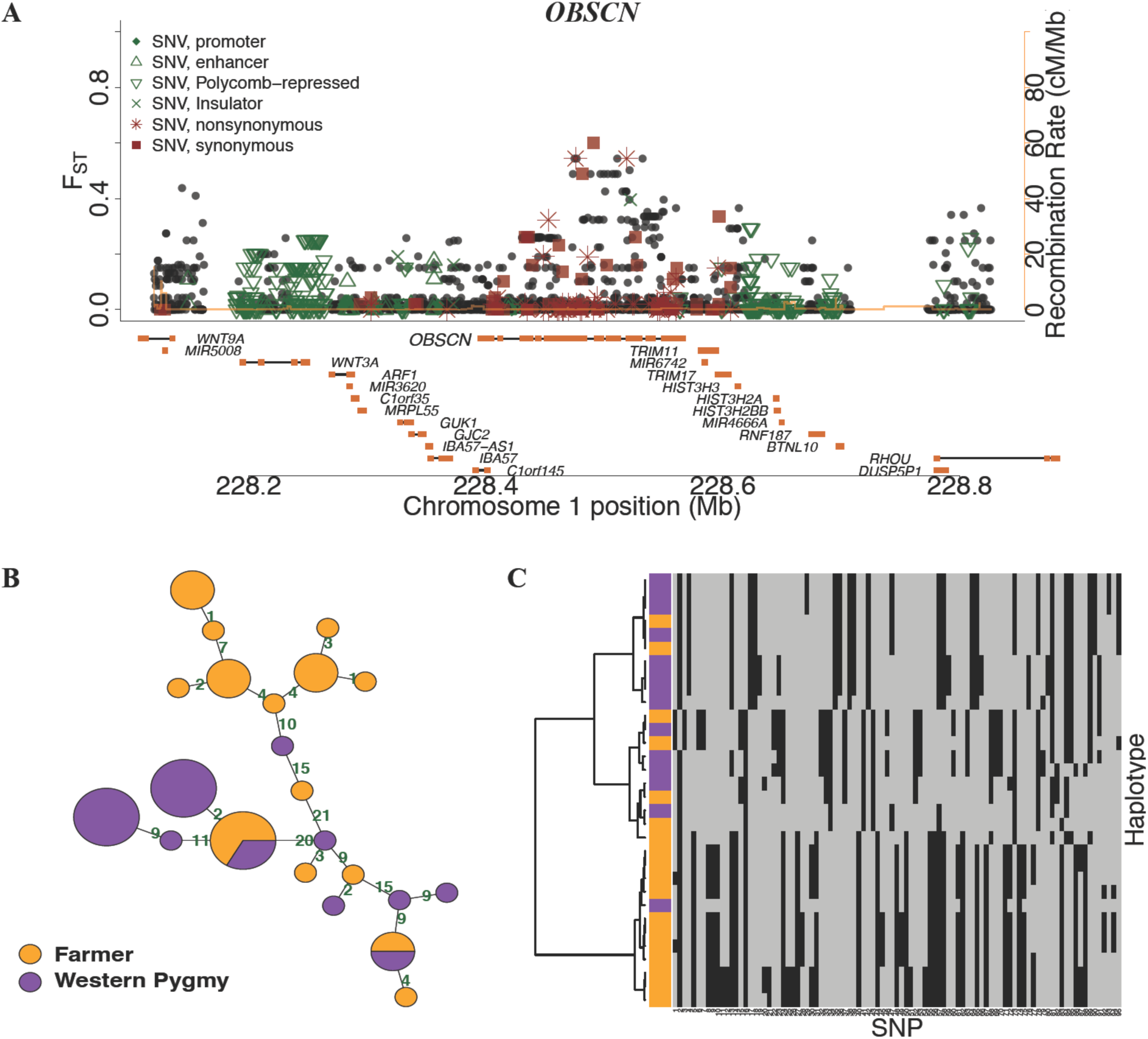
Candidate selection signal near *OBSCN*. (A) As in **Fig. 3**, but for the region chr1:228103665-228842760. (B-C) As in Fig. 3, but for the region chr1: 228.46-228.54 Mb with elevated F_ST_.

Interestingly, Andersen et al. (2012) recently found evidence that variants in *LARGE* might have been positively selected for the resistance of Lassa fever in Western African populations.

Our whole-genome selection scan also identified a variety of genes (**Table S3**) involved in immune function, one of the most common targets of adaptive evolution (Williamson et al. 2007; Barreiro and Quintana-Murci 2009), and in reproduction, which is compatible with the life-history tradeoff hypothesis (Migliano et al. 2007). Other functional categories for genes of potential interest within the top hits of our iHS signals (**Table S3**) include energy metabolism, cell signaling, and neural development.

### Evidence of Local Adaptation in Western African Pygmies: G2D

To complement our iHS scan, we performed a scan using the G2D statistic (Nielsen et al. 2009), which measures how different the local farmer-Pygmy 2-D joint allele frequency spectrum is from the genome-wide spectrum. We found low *P*-value top-hit windows on all 22 chromosomes (**Figure S12**). To identify Pygmy-specific signals of selection, we used the composite likelihood ratio (CLR, Nielsen et al. 2005) statistic, which is essentially the 1-D version of the G2D statistic. Our Pygmy-specific top-hit windows satisfied three conditions for all four simulation sets: 1) they were in the top 0.5% of the *P*-value distribution of the G2D statistic, 2) they were in the top 0.5% of the *P*-value distribution of the Pygmy-specific CLR statistic, and 3) they were not within the top 1% of the *P*-value distribution of the Yoruba-specific CLR statistic. This procedure identified 7 distinct Pygmy-specific candidates (**Table S4**). These candidates do not overlap with those from iHS scan, highlighting the complementarity of the G2D and iHS statistics.

Our top candidate region from the G2D scans (locus: chr6:32968692-33049012; **Figure 5A**; *P*-value=9.90×10^-6^, FDR=0.03) includes three members of the Class II Human Leukocyte Antigen (HLA) gene family, *HLA-DPB1, HLA-DOA,* and *HLA-DPA1*. These genes encode proteins that are expressed on antigen-presenting cells and that present extracellular peptides for T-cell recognition. They thus play a critical role in in initiating the immune response to invading pathogens (Barreiro and Quintana-Murci 2009; O’Brien et al. 2011). The HLA region has a complex genomic architecture with several recombination hotspots (**Figure 5A**; also see Jeffreys et al. 2005). To avoid possible artifacts due to sequencing and genotyping errors, we reanalyzed this region after removing variants violating Hardy-Weinberg Equilibrium, an indicator of possible genotyping errors. 14 out of 1478 SNVs in this region fail the HWE test (cutoff p < 0.05); yet the *P*-value for this region remains the same after their removal. 11 non-synonymous variants were found in this region; of these sites, five are predicted to be deleterious or possibly damaging by SIFT (SIFT score $ 0.05, Kumar et al. 2009) and/or PolyPhen2 (PolyPhen2 score # 0.995, Adzhubei et al. 2010). Haplotype analyses (**Figure 5B-C**) of the region with elevated F_ST_ around the gene *HLA-DPA1* show that while the farmer samples possess two major haplotypes, most of the Pygmy samples belong to a single haplogroup. Because of the existence of several recombination hotspots in this locus, we plotted the diplotypes for this region in our sample to avoid possible biases due to phasing error (**Figure S13**). Consistent with the haplotype analyses, most of the Pygmy samples (5 out of 7) are homozygous for a single diplotype, while the farmers have two diplotypes. We thus hypothesize that a specific immunity-related pressure has driven the evolution of this locus in the Pygmies.

**Figure 5.**
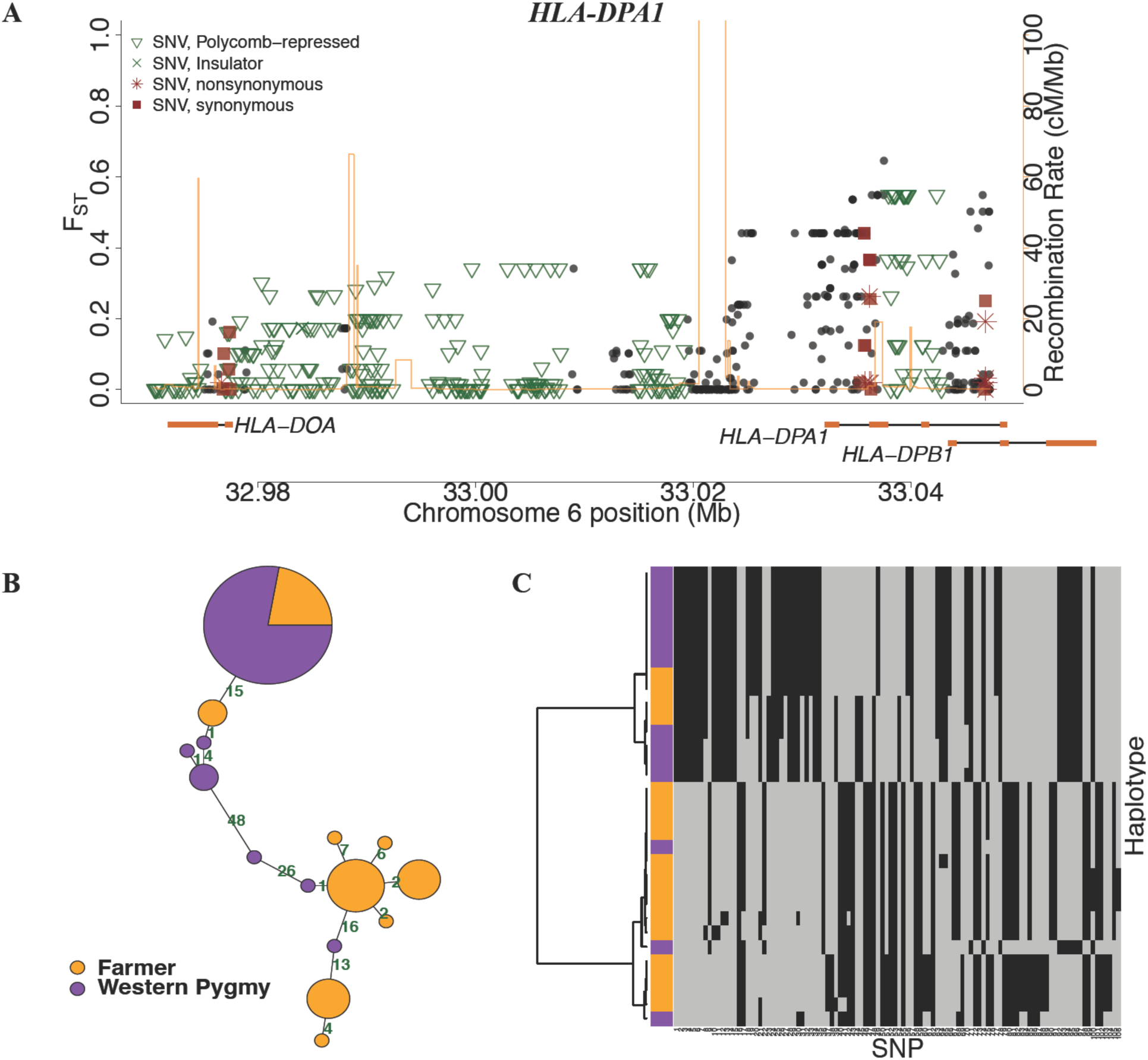
Candidate selection signal near *HLA-DPA1*. (A) As in **Fig. 3**, but for the candidate locus chr6:32968692-33049012. (B-C) As in **Fig. 3**, **but for the** region chr6: 33.03-33.05 Mb with elevated F_ST_.

This scan also identified two candidate regions that contain genes associated with bone synthesis and development. The gene *FLNB* in the first region (locus: chr3:57918877-58055004) encodes filamin B, a multifunctional cytoplasmic protein that plays a critical role in skeletal development. *Flnb* knockout mice are phenotypically similar to individuals with spondylocarpotarsal syndrome as they exhibited short stature and similar skeletal abnormalities (Farrington-Rock et al. 2008). *FLNB* is known to be associated with height in African Pygmies (Jarvis et al. 2012; Lachance et al. 2012) and has also been reported to be associated with osteoporosis in women (Wilson et al. 2009). Although we did not find any amino acid substitution variants in *FLNB* in our sample, we did find many variants with large F_ST_ that may lie in regulatory elements (**Figure S14**). The second region (locus: chr1:179361049-179468857) contains the gene *AXDND1*. Although the function of *AXDND1* is still unclear, a recent multiple-cohort genome-wide association study reported a statistically significant association between this gene and fracture risk. This implies a potential role of *AXDND1* in bone synthesis or other musculoskeletal traits (Medina-Gomez and Rivadeneira 2014).

One of our candidate regions (locus: chr1:183076845-183184161) includes the gene *LAMC1*, which plays a role in reproductive development. *LAMC1* expression increases in bovine, pig, and rabbit basal lamina during follicular development (Irving-Rodgers and Rodgers 2005), and is also expressed in the human ovary (Berkholtz et al. 2006). A recent genome-wide association study reported that polymorphisms in *LAMC1* are associated with an increased risk of premature ovarian failure, which is characterized as the cessation of ovarian function before the age of 40 and could result in amenorrhea and infertility (Pyun et al. 2012). Another interesting candidate region (chr19:12386669-12523799) contains cell signal transmission genes, the *ZNF* genes, which encode proteins with KRAB and zinc-finger domains. Genes in this protein family have been previously shown to be under positive selection in African Americans (Nielsen et al. 2005; Nielsen et al. 2009). It is unclear what phenotype these variants are associated with, but the role of *ZNF442* in transcriptional binding activity suggests *trans*-regulatory evolution might play a role in the adaptation of Pygmies.

### Inference of Polygenic Selection in Western African Pygmies

Many human traits, such as body weight and height and skin pigmentation, are polygenic. Our G2D and iHS scans were designed to detect large changes in allele frequency at single loci, but polygenic selection can result in small changes in allele frequency at multiple loci that involved in a specific biological function or pathway (Pritchard et al. 2010; Berg and Coop 2014). To detect polygenic selection, we used the F_ST_ statistic (Weir and Cockerham’s estimator) to estimate the level of population differentiation for each SNV and compared the F_ST_ distribution of the SNVs of all genes in a given gene set (a specific biological function or pathway) to that in the rest of our genic sequences. We used 1,454 Gene Ontology (GO) gene sets downloaded from the Gene Set Enrichment Analysis (GSEA) project (Subramanian et al. 2005).

Using the Mann Whitney U test, we found 113 gene sets that show significant evidence of having larger F_ST_ values compared to the F_ST_ distributions of the rest of genic sequences (one-sided test, Bonferroni corrected p < 10^-10^). To evaluate our false positive rate, we conducted the same analysis using our four sets of neutral whole genome simulations. Surprisingly, we found as many or more significant gene sets in 72.9% and 48.5% of our simulations under the Model-1 and Model-2, respectively. This suggests that demographic processes and genomic architecture can mimic the signals of polygenic adaptation, and in turn suggests that many of these 113 significant gene sets are false positives.

Only 3 out of the 113 significant gene sets had significant U tests less than 5% of the time in all of our neutral whole genome simulation sets, and we consider these sets as true positives (**Table S5**). Among these three gene sets, there were no overlapping genes, nor did any genes overlap with those identified in our G2D and iHS analysis. The two strongest signals of polygenic selection we detected are both related to immunity. The Gene Ontology (GO) category “Antigen binding” (Bonferroni p-value = 2.31×10^-25^) consists of antigen binding proteins, which interact selectively with an antigen or any substance which is capable of inducing a specific immune response and of reacting with the products of that response (GO:0003823). The GO category “Pattern Recognition Receptor Activity” (Bonferroni p-value = 5.04×10^-14^) includes molecules in the innate immune system that recognize microorganisms through a series of pattern recognition receptors and that are highly conserved in evolution (Dziarski 2003). Although no overlap among candidate regions is observed from the iHS, G2D, and polygenic selection tests, the pervasive selective signals for immunity in all three tests highlight the importance of adaptation against parasites and infectious disease pathogens in Pygmy evolution. The other candidate of polygenic selection is the GO category “G1 phase of mitotic cell cycle” (Bonferroni p-value = 1.75×10^-19^). Although the corresponding phenotype for this group is unknown, accurate transition from G1 phase of the cell cycle is crucial for control of eukaryotic cell proliferation (Bertoli et al. 2013).

## Discussion

### Ancient Divergence and More Recent Gene Flow between African Farmer and Pygmy Populations

Our demographic inference for the farmer (Yoruba) and Western Pygmy hunter-gatherer groups (Baka and Biaka) offers insight into the demographic dynamics of sub-Saharan Africa over the past hundreds of thousands years. The deep divergence time between the ancestors of the agricultural and Pygmy groups we found in Model-1 (~155 kya, 95% C.I.: 139–164 kya) is inconsistent with several recent publications (Patin et al. 2009; Batini et al. 2011; Veeramah et al. 2011), in which 95% confidence intervals of divergence times were estimated to be 25–130 kya (Patin et al. 2009), 51-106 kya (Batini et al 2011), and 10-105 kya (Veeramah et al 2011). In our Model-2, although still relatively old, this divergence time (~90 kya, 95% C.I. 85– 91 kya) is more consistent with those earlier studies. Our PSMC analysis in consistent with old divergence between the ancestors of the two groups (**Figure S2**); but these results must be interpreted carefully because the PSMC does not explicitly model population divergence. In the late Pleistocene, the African continent experienced dramatic climate fluctuations between dry and wet conditions near the end of Marine Isotope Stage 6 (MIS 6, 190-135 kya) and through the whole MIS 5 (75–135 kya) (Blome et al. 2012; Rito et al. 2013). Paleoclimatic and archaeological data suggest that late Pleistocene hominin movements and occupation favored drier intervals, when droughts caused forest defragmentation and open grassland expansion (Blome et al. 2012, Ziegler et al. 2013). Within tropical Central Africa, human populations were relatively buffered from the effects of climate change, so this region may have provided refugia during the glacial MIS 6 and MIS 5 (Blome et al. 2012, Rito et al. 2013, Ziegler et al. 2013). These refugia may have in turn promoted population isolation and differentiation (Blome et al. 2012, Ziegler et al. 2013). Combining these climatic observations with our genetic inferences, we speculate that environmental change and forest fragmentation may have caused the ancestors of Pygmy rainforest dwellers to diverge from the ancestors of agricultural groups within the past 75–190 kya. Although Model-1 is the best-fit model for our data, the deep Pygmy-farmer divergence could be in part due to imperfections in the model. For example, our model does not incorporate archaic introgression, which has been reported recently in Western African Pygmies (Hammer et al. 2011). Such introgression might cause us to overestimate the Pygmy-farmer divergence.

Our inference of asymmetric gene flow in Model-1 is consistent with observed socio-economic contacts and intermarriage practices between Pygmies and farmers (Terashima 1987; Terashima 1998; Bahuchet 2012), and was also observed previously in Patin et al. (2009). However, Patin et al. (2009) had little power to infer gene flow since divergence (95% C.I. covers 0). Our best-fit model, Model-1 involves strongly asymmetric gene flow that did not begin until 40 ky (C.I.: 36–43 ky) after the ancestors of Pygmies and farmers diverged. This timing coincides with the aftermath of most of the large waves of population expansion and technological improvement within Africa, dated 55–75 kya (Henshilwood et al. 2002, Ziegler et al. 2013). This early improvement in hunting tools, stone, and shell ornaments might have increased the carrying capacity of populations associated with this radical technological evolution, and thus might also have promoted contacts between different human groups, including the ancestors of farmers and Pygmy hunter-gatherers.

There are important differences between the approach used here and those used in earlier demographic studies of African Pygmies (Patin et al. 2009, Batini et al. 2011, Veeramah et al. 2011). First, we jointly estimated all parameters simultaneously for a given model, but some previous studies first estimated effective population sizes and then optimized other model parameters given the pre-estimated population sizes (Patin et al. 2009). They thus explored a smaller region of parameter space, potentially biasing their inferences. Second, our inference was based on whole genome sequencing data with a relative small sample size of 16 genomes, while these previous studies all used less than 60 loci, but had much larger samples of >100 individuals. Two of these studies (Patin et al. 2009; Batini et al. 2011) inferred recent population contraction in the Pygmy groups, a process our relatively small sample may not have had power to detect (Robinson et al. 2014).

Our second best-fit model, Model-2, suggests a single pulse of gene flow from farmers to Pygmies ~7 kya (C.I.: 6–7 kya), resulting in a ~68% (C.I.: 67.9%–68.2%) admixture in our Western Pygmy sample. The observation of substantial agriculturalist genetic ancestry in Pygmies has been hypothesized as the consequence of the recent expansion of Bantu/Niger-Kordofanian-speaking farmers from West Africa about 5 kya (Cavalli-Sforza 1986; Tishkoff et al. 2009; Patin et al. 2014). The observed substantial admixture proportion in our Western Pygmy sample is consistent with the recent finding of Verdu et al. (2013), which analyzed autosomal microsatellite data of 800 individuals among 23 Central African Pygmy and non-Pygmy populations and inferred admixture proportions of up to 50-70% in these Pygmy populations. Other studies also found evidence of genetic admixture in African Pygmies although their proportions vary between 0-90% among individual Pygmies (Patin et al. 2009; Tishkoff et al. 2009; Jarvis et al. 2012; Patin et al. 2014). Our inferred time of admixture coincides with the time of Neolithic agricultural development in Africa about 5–10 kya (Phillipson 2005), as well as with the estimated times of agriculturalist expansion for both Bantu-speaking (5.6 kya, 95% C.I.: 3.2–8.2 kya) and Niger-Kodorfanian-speaking (7.3 kya, 95% C.I.: 5.7–9.6 kya) people, reported by Li et al. (2014). Many Pygmies today speak languages adopted from neighboring Bantu-and/or Sudanic-speaking farmer groups, with whom they exchange goods (Bahuchet 2012). Because the social-economic relationship between the two groups can sometimes promote intermarriage (Terashima 1987, Bahuchet 2012), this symbiotic bond may contribute to the observed substantial admixture in the Pygmy groups, especially since the development and expansion of agriculture in Africa.

Our inferred dates are based on a phylogeny-based mutation rate of 2.35×10^-8^ per-site per-generation (Gutenkunst et al. 2009; compatible with Nachman and Crowell 2000) and a generation time of 25 years. Our date estimates would be much older if we used the rate of ~1.2×10^-8^ per-site per-generation estimated by recent pedigree-based whole genome sequence studies (Conrad et al. 2011; Kong et al. 2012). Both approaches to estimating the human mutation rate have limitations, including inaccuracy of the human-chimpanzee divergence time in the phylogenetic approach and false negative mutations in the pedigree sequencing approach (Veeramah and Hammer 2014). We used the phylogenetic estimate because of its history in population genetic inference, but caution is advised when comparing population genetic date estimates with the archeological and fossil record.

### Importance of Prioritizing Selection Candidates Using P-values From Whole Genome Simulations

Our results highlight the importance of using a model-based approach to assess statistical significance in whole-genome selection scans. Genomic scan studies using the tail of an empirical summary statistic distribution (an “outlier” approach) to define a significance cutoff for positive selection have been highly criticized. Non-selective forces, including demography and local genomic architecture, such as variation in mutation and recombination rates (Reich et al. 2002; Drake et al. 2005; Jeffreys et al. 2005; Schaffner et al. 2005; Sainudiin et al. 2007) across loci, can produce signals similar to positive selection (Tajima 1989; Andolfatto and Przeworski 2000; Wall et al. 2002; Jensen et al. 2005; Schaffner et al. 2005; Teshima et al. 2006). For example, we observed that larger G2D scores are associated with higher heterozygosity (**Figure S6**), so candidates determined using an empirical outlier approach might be biased towards regions with higher mutation rates. By matching local mutation rate in our simulations to local heterozygosity in the data, we eliminate this bias (**Figure S7**). Worryingly, the false targets identified by a genomic scan that fails to account for non-selective forces can be misleading because they might still make biological sense a posteriori (Pavlidis et al. 2012).

Many outlier windows in our empirical distributions are not atypical when the underlying demography and genomic architecture are controlled for (**Figure 2, Quadrant I**). Prioritizing selection candidates based on *P*-values also identifies regions whose summary statistic values are modest, yet atypical compared to the same regions in our simulations (**Figure 2, Quadrant III**). Even more striking is the high proportion of GO gene sets that are identified as significant by a Mann-Whitney U test but that are not significant when compared against our neutral simulations that account for demographic history and genomic architecture. Caution is still advised when interpreting our results, however, because no simulation can account for all potential confounding factors (Pavlidis et al. 2012).

### Candidates of adaptation in Western African Pygmy groups

With our high coverage whole-genome sequencing data, we conducted a comprehensive model-based whole-genome scan for natural selection for Western African Pygmies using a series of complementary statistical approaches. Many loci detected by our approach are involved in muscle development, bone synthesis, immunity, reproduction, cell signaling and development, and energy metabolism (see Results).

Of particular interest are several genomic regions that show signatures of selection in African Pygmies that might contribute to short stature. Seven genes known to be associated with bone synthesis were identified by either iHS or G2D analysis. Among them, *FLNB*, *EPHB1*, and *TSPAN5* have been functionally shown to affect body size through gene knockout or knockdown experiments in mice (Iwai et al. 2007; Farrington-Rock et al. 2008; Benson et al. 2012; Zhou et al. 2014), and *FLNB*, *AXDND1*, *ZBTB38*, and *GAREM* have been shown to be associated with human height in multiple populations (Lettre et al. 2008; Weedon et al. 2008; Kim et al. 2012; Wang et al. 2013; Medina-Gomez and Rivadeneira 2014; Wood et al. 2014). *EPHB1* was reported to be genetically associated with height in African Pygmies (Jarvis et al. 2012). Interestingly, although we found no non-synonymous variants in the locus containing *EPHB1*, the Pygmy and farmer populations are each nearly fixed for a single population-specific haplotype (**Figure 3B-C**), an pattern expected under an incomplete selective sweep (Voight et al. 2006, Pickrell et al. 2009, Pritchard et al. 2010). *FLNB* (locus: chr3:57,918,877-58,055,004) is within the locus chr3:45–60Mb that was also previously reported to be associated with height in Pygmies (Jarvis et al. 2012, Lachance et al. 2012). Clinically, nonsense mutations in *FLNB* cause of Spondylocarpotarsal synostosis syndrome (SCT), a recessive disease characterized by short stature and fusions of the vertebrae and carpal and tarsal bones (Krakow et al. 2004). Our observation of many large F_ST_ variants within ENCODE regulatory sequences (**Figure S14**) around this locus suggests that short stature in Western African Pygmies might arise through *cis*-regulatory evolution.

Several studies (e.g. Diamond 1991; Venkataraman et al. 2013) have hypothesized that the ability to quickly climb trees and move in dense forest is a potential adaptation of Pygmy hunter-gatherers. The most highly differentiated SNV rs437129 in our candidate gene *OBSCN*, a myofibrils regulating obscurin gene, is predicted by PolyPhen2 (score = 0.968) to be a functionally important nonsynonymous variant (although not by SIFT; SIFT score = 0.43). Our haplotype analyses suggest that this SNV is associated with population-specific haplotypes in the Pygmies and the farmers (**Figure 4B-C**), although the signal is noisy. In Pygmies, the fixed allele of this SNV is consistent with the ancestral state (PanTro3, Hg19). Under a classic selective sweep model, one might expect a derived beneficial allele to sweep up in frequency, but a nearby ancestral allele could hitchhike with the selected site (Smith and Haigh 1974). However, selection may sometimes favor an ancestral allele that has been segregating in the population (Pritchard et al. 2010). Because accessing essential foods is crucial for hunter-gatherers, mobility-related adaptation to locomotor efficiency amid dense vegetation has been emphasized in several recent studies (Diamond 1991; Bramble and Lieberman 2004; Perry and Dominy 2009). Indeed, Venkataraman et al. (2013) recently presented evidence of a positive correlation between tree climbing ability and muscle fiber length in African Twa and Asian Agta Pygmies compared to neighboring non-tree-climbing farmers. This suggests that tree falls could be a substantial selective pressure, and natural selection might in turn have favored anatomical structures (e.g. muscle fiber length) that promote safe vertical climbing (Venkataraman et al. 2013). A plausible evolutionary explanation for our observed selective signal is that natural selection favors the ancestral haplotype of *OBSCN* possessed in hunter-gatherer Pygmies to adapt specific muscle architecture to locomotor efficiency, while local adaptation outside the forest to an alternative allele or relaxation of selection might promote the observed population differentiation around this locus. The selective signal we found around the gene *OBSCN* could thus be the first genetic evidence that supports the mobility hypothesis.

We employed several complementary statistical tests to detect different modes of adaptation. The haplotype-based iHS test has greatest power for detecting recent (< 30 kya) incomplete sweeps, but the frequency-spectrum-based G2D test is capable of detecting completed and ongoing sweeps that occurred < 300 kya as well as balancing selection (Sabeti et al. 2006; Nielsen et al. 2009). Our gene set enrichment analysis, on the other hand, has little power to detect sweeps but can detect polygenic selection (Daub et al. 2013). It is thus not surprising that there is no overlap among the candidates identified by our different tests. All three tests did, however, detect regions including genes associated with immunity (see Results). In particular, our analyses suggest an ongoing sweep in Pygmies near the immunity gene *HLA-DPA1* (**Figure 5**). The pervasive signals of natural selection on immune function we find are consistent with the view that genes involved in pathogen response are among the most common targets of adaptive evolution (Williamson et al. 2007; Barreiro and Quintana-Murci 2009; Jarvis et al. 2012; Novembre and Han 2012).

We leveraged whole-genome sequence data from African Pygmy and agriculturalist populations to infer their prehistory and search for Pygmy-specific adaptation signals through a carefully designed computational and statistical framework. In doing so, we accounted for many potentially confounding factors, including demography and mutation and recombination rate heterogeneity. Future work may be needed to account for additional confounding factors, but we believe the framework presented here offers great promise for shedding light on the complex demographic and adaptive history of human populations.

## Materials and Methods

### Whole genome sequencing data and data quality assurance

Our Biaka Pygmy (N=4) DNA samples were obtained from publicly available cell lines administrated by the Centre d’Etude du Polymorphism Human Genome Diversity Panel (Li et al. 2008). Details regarding the Baka Pygmies (N=3) samples are in Lachance et al. (2012). Whole-genome sequencing data for the unrelated Yoruba farmers (N=9) were downloaded from the CGI data repository (Coriell sample IDs: NA18501, NA18502, NA18504, NA18505, NA18508, NA18517, NA19129, NA19238, and NA19239). The median coverage across the samples was 60.5X (s.d. = 8.54X). Genome assembly and variant calling were done using the standard CGI Assembly Pipeline 1.10, CGA Tools 1.4, and NCBI Human Reference Genome build 37. Before any quality control filters, 13,276,198 autosomal single nucleotide variants (SNVs) were called in our samples. Unless mentioned otherwise, we analyzed only variants that were 1) fully called across all samples, 2) not in any known or called indels, 3) not in any known or called copy number variants, 4) not in any known segmental duplication regions, and 5) aligned against chimpanzee (PanTro3, Hg19). Databases that used for steps 3, 4, and 5 were downloaded from UCSC Genome Browser in May 2013. We used Hg19 coordinates, using the UCSC Genome Browser liftOver program if necessary. After filtering, our data consist of 10,865,288 SNVs.

### Estimation of demographic parameters using ∂a∂i

We used the demographic inference tool *∂*a*∂*i (Gutenkunst et al. 2009) to build and fit our demographic. In short, *∂*a*∂*i is a forward time simulator of allele frequency spectrum (AFS) based on a diffusion approximation (Kimura 1964). To ensure genotype quality for demographic inference, SNVs were removed if they overlapped with any known repetitive genomic regions based on the UCSC Genome Browser databases, Self Chain (if sequence identity > 0.9) and RepeatMasker. Sites within genes or 1,000 flanking base pairs were excluded to minimize possible effects of natural selection. Coordinates of genes were from the RefSeq genes database, downloaded from the UCSC Genome Browser in May 2013. We used the remaining 1,575,394 SNVs from a total of 325,957,426 non-genic base pairs to build an unfolded AFS, polarized via human-chimpanzee alignment (PanTro3, Hg19). We used the *∂*a*∂*i implementation of a context-dependent substitution model to statistically correct the unfolded AFS to mitigate possible biases due to ancestral state misidentification (Hernandez et al. 2007). To estimate demographic parameters, the derivative-based BFGS algorithm was used to optimize the composite log-likelihood. Confidence intervals were estimated fitting 100 non-parametric bootstraps of the non-genic data. We converted parameters from the population-genetic units to physical units using a phylogenetic-based mutation rate of 2.35×10^-8^ per-base per-generation (Nachman and Crowell 2000; Gutenkunst et al. 2009) and a generation time of 25 years.

### Assessment of demographic model

The composite likelihood *∂*a*∂*i calculates is not the full likelihood due to the linkage. To minimize linkage in our model selection analysis, we thinned our data by choosing variants at least 0.01 cM apart and re-fit the candidate models to the resulting sub-dataset. We then calculated AIC (Akaike 1974) and BIC (Schwarz 1978) as -2log*L* + *kP,* where *L* is the likelihood function, *k* is the number of parameters in the model, and *P* is 2 for AIC and log(*n*) for BIC, where n is the sample size. In our comparisons of real and simulated LD decay, we estimated LD between pairs of variants by their correlation coefficient (*r*^*2*^) using a genotype code (0, 1, or 2 reference alleles). We performed our PSMC analysis with v0.6.3 (Li & Durbin 2011), using the parameters suggested by the authors: psmc -N25 -t15 -r5 -p “4+25*2+4+6”. To assess variation in the inferred PSMC curves, we analyzed 100 non-parametric bootstraps using the utility provided in the PSMC software.

### Coalescent whole-genome simulations

We used MaCS (Chen et al. 2009) for our coalescent simulations, because of its ability to efficiently perform whole genome simulations with recombination. To avoid potential underestimation of recombination rates, we removed the first 5 Mb on each chromosome as suggested by the creators of the African American recombination map (Hinch et al. 2011). For consistency, we also did this for the HapMap map (Frazer et al. 2007). To model mutational heterogeneity, carried out a 3-step procedure. First, we divided the genome into 25,000 bp windows and estimated the population genetic mutation parameter 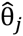 using *∂*a*∂*i given a demographic model. Second, we performed each MaCS using a mutation parameter 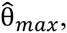 the largest “ estimated among all of the windows. Third, for each window we adjusted its mutation rate by dropping a proportion 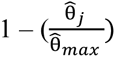 of the simulated variants. All simulations presented here model the effects of demography, recombination heterogeneity, and mutation heterogeneity. For our simulations, we excluded the regions that were excluded in the real data due to our quality control criteria.

### Scan for signals of selective sweeps

All test statistics were calculated using pre-defined sliding windows of 500 SNVs, with a step size of 100 SNVs. Windows longer than 1 Mb were dropped to avoid complex genomic regions, such as centromeres or large structure variants. To maximize statistical power and focus on signals of selection in Western Pygmies generally, for all our tests we combined samples from the two Pygmy populations because they are so recently diverged. We calculated statistical significance of each window using our whole-genome coalescent simulations under the best-fit demographic models. To account uncertainty in parameter, we drew 1,000 parameter sets from the confidence intervals from each model, assuming that they had a multivariate normal distribution. The per-window *P*-value was defined as the fraction of simulations with statistic values greater or equal to the observed value of the same window in the real data. Candidates for each neutrality test were defined as the top 0.5% of the corresponding *P*-value distribution. We then ran 100,000 additionally local simulations for each candidate window to obtain a finer *P*-value resolution. We estimated false discovery rates using the method of Williamson et al. (2006).

The G2D test searches for evidence of hard selective sweeps by comparing the spatial distribution of the frequency spectrum of a locus to the expected one based on the whole genome (Nielsen et al. 2009). This statistic is defined as

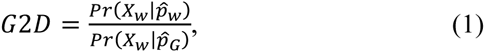

where *X*_*w*_ is the SNV data in the window *w* and 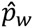 and 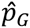 are the estimated joint allele frequency spectra using the SNV data in the window *w* and whole genome, respectively. The G2D statistic is calculated on the unfolded frequency spectrum based on the human-chimpanzee alignment (PanTro3, Hg19). Possible biases due to ancestral state misidentification are accounted for in the inference of the demographic models and, therefore, in our simulations when assessing the statistical significance.

To detect regions undergoing a recent partial sweep, we used the integrated haplotype score (iHS, Voight et al. 2006) implemented in the software selscan (Szpiech and Hernandez 2014). Haplotype phasing was done using BEAGLE v3.1.1 (Browning and Browning 2007). To enhance phasing accuracy, we included additional public genotype data: a Bakola and a Bedzen genome (CGI Assembly Pipeline 1.10, CGA Tools 1.4) from Lachance et al. (2012), 16 Biaka Pygmies genotyped by the Human Genome Diversity Project (HGDP, Li et al. 2008, Illumina 650 K), and 10 Baka and 10 Bakola Pygmies genotyped by the Hammer lab of the University of Arizona (Affymatrix Axiom 500K). The 9 Yoruba genomes were phased separately using the same approach, with an additional 4 Luhya genomes from the CGI public data repository, genotype data of 81 Yoruba and 86 Luhya samples from the 1000 Genomes Project and 21 Yoruba and 10 Luhya samples from the HGDP. All positions were converted into Hg19 coordinates using UCSC LiftOver utility if necessary. We filtered out duplicated samples and/or related individuals using the identical-by-descent operation in the software package PLINK (Purcell et al. 2007). To account for possible biases in the downstream iHS analysis due to phasing errors, haplotype phase was estimated using the same procedure used for the real data. iHS was calculated with the default parameters in selscan, and standardized following Voight et al. (2006). For both real and simulation data, the strength of selection signal in each window was quantified by the proportion of SNVs with |iHS| > 2 (Voight et al. 2006).

### Haplotype and diplotype analyses

Hierarchical clustering for both haplotype and diplotype data was performed using the R function “hclust” in the stats package (R Development Core Team, 2012). We used the R package pegas (v.0.6, Paradis 2010) to plot haplotype network, using pairwise nucleotide differences as the distance matrix.

### Inference of polygenic selection

We downloaded 1,454 Gene Ontology gene sets from the Gene Set Enrichment Analysis (GSEA) project at the Broad Institute in January 2014 (Subramanian et al. 2005), discarding 13 gene sets that shared more than 90% of their genes with another set. One-sided (alternative distribution is greater than the null) Mann-Whitney U tests were performed in R (R Development Core Team, 2012). In our simulations, the genic F_ST_ distributions were obtained by calculating F_ST_ for all SNVs within the same genomic regions that are defined as genes in the real data (RefSeq, downloaded from USCS Genome Browser in May 2013). The likelihood of a gene set being significant was calculated as

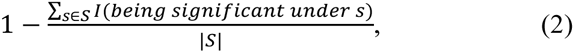

where |*S*| is the total number of whole genome simulations, *s* is a given whole genome simulation, and *I* is an indicator function of being significant under *s*.

### ENCODE regulatory elements

We downloaded the ENCODE database (wgEncodeBroadHmmHsmmHMM) using the UCSC Genome Browser in February 2014. We used the five most reliable functional categories: Active Promoter (state 1), Strong Enhancer (states 4 and 5), Insulator (state 8), and Polycomb-repressed (state 12). This yielded 134,769 regulatory elements.

## Data Accessibility

Our Biaka genomic data will be available in the dbSNP database (http://www.ncbi.nlm.nih.gov/projects/SNP/). The Baka genomic data are in dbSNP with submitter batch IDs: Lachance2012Cell_snp, Lachance2012Cell_deletion, Lachance2012Cell_insertion, Lachance2012Cell_complex_substitution. The Yoruba genomic data are available at the CGI public data repository (http://www.completegenomics.com/public-data/).

## Acknowledgements

Support for this work was provided by the National Institutes of Health to J.D.W and M.F.H (R01 HG005226). P.H. and R.N.G. were supported by National Science Foundation (NSF) grant DEB-1146074. S.A.T was supported by National Institute of Health (NIH) grants 1R01GM113657-01 and 8DP1ES022577-04. J.L. is grateful for support from NIH NRSA postdoctoral fellowship F32HG006648.

## References

Ackermann MA, Shriver M, Perry NA, Hu L-YR, Kontrogianni-Konstantopoulos A. 2014. Obscurins: Goliaths and Davids Take over Non-Muscle Tissues. PloS one 9(2): e88162.

Adzhubei IA, Schmidt S, Peshkin L, Ramensky VE, Gerasimova A, Bork P, Kondrashov AS, Sunyaev SR. 2010. A method and server for predicting damaging missense mutations. Nature methods 7(4): 248–249.

Akaike H. 1974. A new look at the statistical model identification. Automatic Control, IEEE Transactions on 19(6): 716–723.

Akey JM. 2009. Constructing genomic maps of positive selection in humans: where do we go from here? Genome research 19(5): 711–722.

Andersen KG, Shylakhter I, Tabrizi S, Grossman SR, Happi CT, Sabeti PC. 2012. Genome-wide scans provide evidence for positive selection of genes implicated in Lassa fever. Philosophical transactions of the Royal Society of London Series B, Biological sciences 367(1590): 868–877.

Andolfatto P, Przeworski M. 2000. A genome-wide departure from the standard neutral model in natural populations of Drosophila. Genetics 156(1): 257–268.

Bahuchet S. 2012. Changing language, remaining pygmy. Hum Biol 84(1): 11–43.

Barreiro LB, Quintana-Murci L. 2009. From evolutionary genetics to human immunology: how selection shapes host defence genes. Nat Rev Genet 11(1): 17–30.

Batini C, Lopes J, Behar DM, Calafell F, Jorde LB, Van der Veen L, Quintana-Murci L, Spedini G, Destro-Bisol G, Comas D. 2011. Insights into the demographic history of African Pygmies from complete mitochondrial genomes. Molecular biology and evolution 28(2): 1099–1110.

Baumann G, Shaw MA, Merimee TJ. 1989. Low levels of high-affinity growth hormone-binding protein in African pygmies. New England Journal of Medicine 320(26): 1705–1709.

Benson MD, Opperman LA, Westerlund J, Fernandez CR, San Miguel S Henkemeyer M, Chenaux G. 2012. Ephrin-B stimulation of calvarial bone formation. Developmental Dynamics 241(12): 1901–1910.

Berg JJ, Coop G. 2014. A population genetic signal of polygenic adaptation. Plos Genet 10(8): e1004412.

Berkholtz CB, Shea LD, Woodruff TK. 2006. Extracellular matrix functions in follicle maturation. In Seminars in reproductive medicine, Vol 24, p. 262. NIH Public Access.

Bertoli C, Skotheim JM, de Bruin RA. 2013. Control of cell cycle transcription during G1 and S phases. Nature Reviews Molecular Cell Biology 14(8): 518–528.

Blench R. 2006. Archaeology, language, and the African past. Rowman Altamira.

Blome MW, Cohen AS, Tryon CA, Brooks AS, Russell J. 2012. The environmental context for the origins of modern human diversity: a synthesis of regional variability in African climate 150,000-30,000 years ago. Journal of human evolution 62(5): 563–592.

Bramble DM, Lieberman DE. 2004. Endurance running and the evolution of Homo. Nature 432(7015): 345–352.

Browning SR, Browning BL. 2007. Rapid and accurate haplotype phasing and missing-data inference for whole-genome association studies by use of localized haplotype clustering. The American Journal of Human Genetics 81(5): 1084–1097.

Cavalli-Sforza LL. 1986. African pygmies. Academic Pr.

Cavalli-Sforza LL, Menozzi P, Piazza A. 1994. The history and geography of human genes. Princeton university press.

Chen GK, Marjoram P, Wall JD. 2009. Fast and flexible simulation of DNA sequence data. Genome research 19(1): 136–142.

Conrad DF, Keebler JEM, DePristo MA, Lindsay SJ, Zhang YJ, Casals F, Idaghdour Y, Hartl CL, Torroja C, Garimella KV et al. 2011. Variation in genome-wide mutation rates within and between human families. Nature genetics 43(7): 712–U137.

Dávila N, Shea BT, Omoto K, Mercado M, Misawa S, Baumann G. 2002. Growth hormone binding protein, insulin-like growth factor-I and short stature in two pygmy populations from the Philippines. Journal of Pediatric Endocrinology and Metabolism 15(3): 269–276.

Diamond JM. 1991. Anthropology. Why are pygmies small? Nature 354(6349): 111–112.

Diaz F, Thomas CK, Garcia S, Hernandez D, Moraes CT. 2005. Mice lacking COX10 in skeletal muscle recapitulate the phenotype of progressive mitochondrial myopathies associated with cytochrome c oxidase deficiency. Human molecular genetics 14(18): 2737–2748.

Drake JA, Bird C, Nemesh J, Thomas DJ, Newton-Cheh C, Reymond A, Excoffier L, Attar H, Antonarakis SE, Dermitzakis ET. 2005. Conserved noncoding sequences are selectively constrained and not mutation cold spots. Nature genetics 38(2): 223–227.

Drmanac R, Sparks AB, Callow MJ, Halpern AL, Burns NL, Kermani BG, Carnevali P, Nazarenko I, Nilsen GB, Yeung G. 2010. Human genome sequencing using unchained base reads on self-assembling DNA nanoarrays. Science 327(5961): 78–81.

Dziarski R. 2003. Recognition of bacterial peptidoglycan by the innate immune system. Cellular and Molecular Life Sciences CMLS 60(9): 1793–1804.

Farrington-Rock C, Kirilova V, Dillard-Telm L, Borowsky AD, Chalk S, Rock MJ, Cohn DH, Krakow D. 2008. Disruption of the Flnb gene in mice phenocopies the human disease spondylocarpotarsal synostosis syndrome. Human molecular genetics 17(5): 631–641.

Frazer KA, Ballinger DG, Cox DR, Hinds DA, Stuve LL, Gibbs RA, Belmont JW, Boudreau A, Hardenbol P, Leal SM. 2007. A second generation human haplotype map of over 3.1 million SNPs. Nature 449(7164): 851–861.

Gerstein MB, Kundaje A, Hariharan M, Landt SG, Yan K-K, Cheng C, Mu XJ, Khurana E, Rozowsky J, Alexander R. 2012. Architecture of the human regulatory network derived from ENCODE data. Nature 489(7414): 91–100.

Gutenkunst RN, Hernandez RD, Williamson SH, Bustamante CD. 2009. Inferring the joint demographic history of multiple populations from multidimensional SNP frequency data. Plos Genet 5(10): e1000695.

Hammer MF, Woerner AE, Mendez FL, Watkins JC, Wall JD. 2011. Genetic evidence for archaic admixture in Africa. Proc Natl Acad Sci U S A 108(37): 15123–15128.

Henshilwood CS, d’Errico F, Yates R, Jacobs Z, Tribolo C, Duller GA, Mercier N, Sealy JC, Valladas H, Watts I. 2002. Emergence of modern human behavior: Middle Stone Age engravings from South Africa. Science 295(5558): 1278–1280.

Hernandez RD, Williamson SH, Bustamante CD. 2007. Context dependence, ancestral misidentification, and spurious signatures of natural selection. Molecular biology and evolution 24(8): 1792–1800.

Herráez DL, Bauchet M, Tang K, Theunert C, Pugach I, Li J, Nandineni MR, Gross A, Scholz M, Stoneking M. 2009. Genetic variation and recent positive selection in worldwide human populations: evidence from nearly 1 million SNPs. PloS one 4(11): e7888.

Hinch AG, Tandon A, Patterson N, Song Y, Rohland N, Palmer CD, Chen GK, Wang K, Buxbaum SG, Akylbekova EL. 2011. The landscape of recombination in African Americans. Nature 476(7359): 170–175.

Irving-Rodgers HF, Rodgers RJ. 2005. Extracellular matrix in ovarian follicular development and disease. Cell and tissue research 322(1): 89–98.

Iwai K, Ishii M, Ohshima S, Miyatake K, Saeki Y. 2007. Expression and function of transmembrane-4 superfamily (tetraspanin) proteins in osteoclasts: reciprocal roles of Tspan-5 and NET-6 during osteoclastogenesis. Allergol Int 56(4): 457–463.

Jarvis JP, Scheinfeldt LB, Soi S, Lambert C, Omberg L, Ferwerda B, Froment A, Bodo J-M, Beggs W, Hoffman G. 2012. Patterns of ancestry, signatures of natural selection, and genetic association with stature in Western African pygmies. Plos Genet 8(4): e1002641.

Jeffreys AJ, Neumann R, Panayi M, Myers S, Donnelly P. 2005. Human recombination hot spots hidden in regions of strong marker association. Nature genetics 37(6): 601–606.

Jensen JD, Kim Y, DuMont VB, Aquadro CF, Bustamante CD. 2005. Distinguishing between selective sweeps and demography using DNA polymorphism data. Genetics 170(3): 1401–1410.

Kim J-J, Park Y-M, Baik K-H, Choi H-Y, Yang G-S, Koh I, Hwang J-A, Lee J, Lee Y-S, Rhee H. 2012. Exome sequencing and subsequent association studies identify five amino acid-altering variants influencing human height. Human genetics 131(3): 471–478.

Kimura M. 1964. Diffusion models in population genetics. Journal of Applied Probability 1(2): 177–232.

Kong A, Frigge ML, Masson G, Besenbacher S, Sulem P, Magnusson G, Gudjonsson SA, Sigurdsson A, Jonasdottir A, Jonasdottir A et al. 2012. Rate of de novo mutations and the importance of father’s age to disease risk. Nature 488(7412): 471–475.

Kumar P, Henikoff S, Ng PC. 2009. Predicting the effects of coding non-synonymous variants on protein function using the SIFT algorithm. Nature protocols 4(7): 1073–1081.

Lachance J, Vernot B, Elbers CC, Ferwerda B, Froment A, Bodo J-M, Lema G, Fu W, Nyambo TB, Rebbeck TR. 2012. Evolutionary history and adaptation from high-coverage whole-genome sequences of diverse African hunter-gatherers. Cell 150(3): 457–469.

Lettre G, Jackson AU, Gieger C, Schumacher FR, Berndt SI, Sanna S, Eyheramendy S, Voight BF, Butler JL, Guiducci C. 2008. Identification of ten loci associated with height highlights new biological pathways in human growth. Nature genetics 40(5): 584–591.

Li H, Durbin R. 2011. Inference of human population history from individual whole-genome sequences. Nature 475(7357): 493–496.

Li JZ, Absher DM, Tang H, Southwick AM, Casto AM, Ramachandran S, Cann HM, Barsh GS, Feldman M, Cavalli-Sforza LL. 2008. Worldwide human relationships inferred from genome-wide patterns of variation. science 319(5866): 1100–1104.

Li S, Schlebusch C, Jakobsson M. 2014. Genetic variation reveals large-scale population expansion and migration during the expansion of Bantu-speaking peoples. Proceedings of the Royal Society B: Biological Sciences 281(1793): 20141448.

Longman C, Brockington M, Torelli S, Jimenez-Mallebrera C, Kennedy C, Khalil N, Feng L, Saran RK, Voit T, Merlini L. 2003. Mutations in the human LARGE gene cause MDC1D, a novel form of congenital muscular dystrophy with severe mental retardation and abnormal glycosylation of α-dystroglycan. Human molecular genetics 12(21): 2853–2861.

Medina-Gomez C, Rivadeneira F. 2014. Update on the Genetic Basis of Disorders of the Musculoskeletal System (ECTS 2013). IBMS BoneKEy 11.

Mendizabal I, Marigorta UM, Lao O, Comas D. 2012. Adaptive evolution of loci covarying with the human African Pygmy phenotype. Human genetics 131(8): 1305–1317.

Migliano AB, Romero IG, Metspalu M, Leavesley M, Pagani L, Antao T, Huang DW, Sherman BT, Siddle K, Scholes C et al. 2013. Evolution of the Pygmy Phenotype: Evidence of Positive Selection from Genome-wide Scans in African, Asian, and Melanesian Pygmies. Hum Biol 85(1-3): 251–284.

Migliano AB, Vinicius L, Lahr MM. 2007. Life history trade-offs explain the evolution of human pygmies. Proceedings of the National Academy of Sciences 104(51): 20216–20219.

Nachman MW, Crowell SL. 2000. Estimate of the mutation rate per nucleotide in humans. Genetics 156(1): 297–304.

Nielsen R, Hubisz MJ, Hellmann I, Torgerson D, Andrés AM, Albrechtsen A, Gutenkunst R, Adams MD, Cargill M, Boyko A. 2009. Darwinian and demographic forces affecting human protein coding genes. Genome research 19(5): 838–849.

Nielsen R, Williamson S, Kim Y, Hubisz MJ, Clark AG, Bustamante C. 2005. Genomic scans for selective sweeps using SNP data. Genome research 15(11): 1566–1575.

Novembre J, Han E. 2012. Human population structure and the adaptive response to pathogen-induced selection pressures. Philosophical transactions of the Royal Society of London Series B, Biological sciences 367(1590): 878–886.

O’Brien T, Kohaar I, Pfeiffer R, Maeder D, Yeager M, Schadt E, Prokunina-Olsson L. 2011. Risk alleles for chronic hepatitis B are associated with decreased mRNA expression of HLA-DPA1 and HLA-DPB1 in normal human liver. Genes and immunity 12(6): 428–433.

Ohenjo No, Willis R, Jackson D, Nettleton C, Good K, Mugarura B. 2006. Health of Indigenous people in Africa. The Lancet 367(9526): 1937–1946.

Paradis E. 2010. pegas: an R package for population genetics with an integrated–modular approach. Bioinformatics 26(3): 419–420.

Patin E, Laval G, Barreiro LB, Salas A, Semino O, Santachiara-Benerecetti S, Kidd KK, Kidd JR, Van der Veen L, Hombert J-M. 2009. Inferring the demographic history of African farmers and Pygmy hunter–gatherers using a multilocus resequencing data set. Plos Genet 5(4): e1000448.

Patin E, Siddle KJ, Laval G, Quach H, Harmant C, Becker N, Froment A, Régnault B, Lemée L, Gravel S. 2014. The impact of agricultural emergence on the genetic history of African rainforest hunter-gatherers and agriculturalists. Nature communications 5.

Pavlidis P, Jensen JD, Stephan W, Stamatakis A. 2012. A critical assessment of storytelling: gene ontology categories and the importance of validating genomic scans. Molecular biology and evolution 29(10): 3237–3248.

Perry GH, Dominy NJ. 2009. Evolution of the human pygmy phenotype. Trends in ecology & evolution 24(4): 218–225.

Perry GH, Foll M, Grenier JC, Patin E, Nedelec Y, Pacis A, Barakatt M, Gravel S, Zhou X, Nsobya SL et al. 2014. Adaptive, convergent origins of the pygmy phenotype in African rainforest hunter-gatherers. Proc Natl Acad Sci U S A 111(35): E3596–3603.

Phillipson DW. 2005. African archaeology. Cambridge University Press.

Pickrell JK, Coop G, Novembre J, Kudaravalli S, Li JZ, Absher D, Srinivasan BS, Barsh GS, Myers RM, Feldman MW. 2009. Signals of recent positive selection in a worldwide sample of human populations. Genome research 19(5): 826–837.

Pritchard JK, Pickrell JK, Coop G. 2010. The genetics of human adaptation: hard sweeps, soft sweeps, and polygenic adaptation. Current Biology 20(4): R208–R215.

Purcell S, Neale B, Todd-Brown K, Thomas L, Ferreira MA, Bender D, Maller J, Sklar P, De Bakker PI, Daly MJ. 2007. PLINK: a tool set for whole-genome association and population-based linkage analyses. The American Journal of Human Genetics 81(3): 559–575.

Pyun J-A, Cha DH, Kwack K. 2012. LAMC1 gene is associated with premature ovarian failure. Maturitas 71(4): 402–406.

Reich DE, Schaffner SF, Daly MJ, McVean G, Mullikin JC, Higgins JM, Richter DJ, Lander ES, Altshuler D. 2002. Human genome sequence variation and the influence of gene history, mutation and recombination. Nature genetics 32(1): 135–142.

Rito T, Richards MB, Fernandes V, Alshamali F, Cerny V, Pereira L, Soares P. 2013. The first modern human dispersals across Africa. PloS one 8(11): e80031.

Robinson JD, Coffman AJ, Hickerson MJ, Gutenkunst RN. 2014. Sampling strategies for frequency spectrum-based population genomic inference. BMC evolutionary biology 14(1): 254.

Sabeti P, Schaffner S, Fry B, Lohmueller J, Varilly P, Shamovsky O, Palma A, Mikkelsen T, Altshuler D, Lander E. 2006. Positive natural selection in the human lineage. science 312(5780): 1614–1620.

Sainudiin R, Clark AG, Durrett RT. 2007. Simple models of genomic variation in human SNP density. BMC genomics 8(1): 146.

Schaffner SF, Foo C, Gabriel S, Reich D, Daly MJ, Altshuler D. 2005. Calibrating a coalescent simulation of human genome sequence variation. Genome research 15(11): 1576–1583.

Scheinfeldt LB, Soi S, Tishkoff SA. 2010. Colloquium paper: working toward a synthesis of archaeological, linguistic, and genetic data for inferring African population history. Proceedings of the National Academy of Sciences of the United States of America 107 Suppl 2: 8931–8938.

Schwarz G. 1978. Estimating the dimension of a model. The annals of statistics 6(2): 461–464.

Shea BT, Bailey RC. 1996. Allometry and adaptation of body proportions and stature in African pygmies. American journal of physical anthropology 100(3): 311–340.

Smith JM, Haigh J. 1974. The hitch-hiking effect of a favourable gene. Genetical research 23(01): 23–35.

Subramanian A, Tamayo P, Mootha VK, Mukherjee S, Ebert BL, Gillette MA, Paulovich A, Pomeroy SL, Golub TR, Lander ES. 2005. Gene set enrichment analysis: a knowledge-based approach for interpreting genome-wide expression profiles. P Natl Acad Sci USA 102(43): 15545–15550.

Szpiech ZA, Hernandez RD. 2014. selscan: An Efficient Multithreaded Program to Perform EHH-Based Scans for Positive Selection. Molecular biology and evolution 31(10): 2824–2827.

Tajima F. 1989. Statistical method for testing the neutral mutation hypothesis by DNA polymorphism. Genetics 123(3): 585–595.

Team RDC. 2012. R: A language and environment for statistical computing.

Terashima H. 1987. Why Efe girls marry farmers?: Socio-ecological backgrounds of inter-ethnic marriage in the Ituri Forest of central Africa. African study monographs Supplementary issue 6: 65–83.

Terashima H. 1998. Honey and holidays: The interactions mediated by honey between Efe hunter-gatherers and Lese farmers in the Ituri forest. African study monographs Supplementary issue 25: 123–134.

Teshima KM, Coop G, Przeworski M. 2006. How reliable are empirical genomic scans for selective sweeps? Genome research 16(6): 702–712.

Tishkoff SA, Reed FA, Friedlaender FR, Ehret C, Ranciaro A, Froment A, Hirbo JB, Awomoyi AA, Bodo J-M, Doumbo O. 2009. The genetic structure and history of Africans and African Americans. Science 324(5930): 1035–1044.

Veeramah KR, Hammer MF. 2014. The impact of whole-genome sequencing on the reconstruction of human population history APPLICATIONS OF NEXT-GENERATION SEQUENCING. Nat Rev Genet 15(3): 149–162.

Veeramah KR, Wegmann D, Woerner A, Mendez FL, Watkins JC, Destro-Bisol G, Soodyall H, Louie L, Hammer MF. 2011. An early divergence of KhoeSan ancestors from those of other modern humans is supported by an ABC-based analysis of autosomal resequencing data. Molecular biology and evolution: msr212.

Venkataraman VV, Kraft TS, Dominy NJ. 2013. Tree climbing and human evolution. Proceedings of the National Academy of Sciences 110(4): 1237–1242.

Verdu P, Becker NS, Froment A, Georges M, Grugni V, Quintana-Murci L, Hombert J-M, Van der Veen L, Le Bomin S, Bahuchet S. 2013. Sociocultural behavior, sex-biased admixture, and effective population sizes in Central African Pygmies and non-Pygmies. Molecular biology and evolution 30(4): 918–937.

Voight BF, Kudaravalli S, Wen X, Pritchard JK. 2006. A map of recent positive selection in the human genome. PLoS biology 4(3): e72.

Wall JD, Andolfatto P, Przeworski M. 2002. Testing models of selection and demography in Drosophila simulans. Genetics 162(1): 203–216.

Wang Y, Wang Zm, Teng Yc, Shi Jx, Wang Hf, Yuan Wt, Chu X, Wang Df, Wang W, Huang W. 2013. An SNP of the ZBTB38 gene is associated with idiopathic short stature in the Chinese Han population. Clinical endocrinology 79(3): 402–408.

Weedon MN, Lango H, Lindgren CM, Wallace C, Evans DM, Mangino M, Freathy RM, Perry JR, Stevens S, Hall AS. 2008. Genome-wide association analysis identifies 20 loci that influence adult height. Nature genetics 40(5): 575–583.

Williamson SH, Hubisz MJ, Clark AG, Payseur BA, Bustamante CD, Nielsen R. 2007. Localizing recent adaptive evolution in the human genome. Plos Genet 3(6): e90.

Wilson SG, Jones MR, Mullin BH, Dick IM, Richards JB, Pastinen TM, Grundberg E, Ljunggren Ö, Surdulescu GL, Dudbridge F. 2009. Common sequence variation in FLNB regulates bone structure in women in the general population and FLNB mRNA expression in osteoblasts in vitro. Journal of Bone and Mineral Research 24(12): 1989–1997.

Wood AR, Esko T, Yang J, Vedantam S, Pers TH, Gustafsson S, Chu AY, Estrada K, Luan Ja, Kutalik Z. 2014. Defining the role of common variation in the genomic and biological architecture of adult human height. Nature genetics.

Young P, Ehler E, Gautel M. 2001. Obscurin, a giant sarcomeric Rho guanine nucleotide exchange factor protein involved in sarcomere assembly. The Journal of cell biology 154(1): 123–136.

Zhang Z, Xia W, He J, Zhang Z, Ke Y, Yue H, Wang C, Zhang H, Gu J, Hu W. 2012. Exome Sequencing Identifies *SLCO2A1* Mutations as a Cause of Primary Hypertrophic Osteoarthropathy. The American Journal of Human Genetics 90(1): 125–132.

Zhou J, Fujiwara T, Ye S, Li X, Zhao H. 2014. Downregulation of Notch Modulators, Tetraspanin 5 and 10, Inhibits Osteoclastogenesis in Vitro. Calcified tissue international 95(3): 209–217.

